## Supplementary figure S1 for "Prefrontal attentional saccades explore space rhythmically"

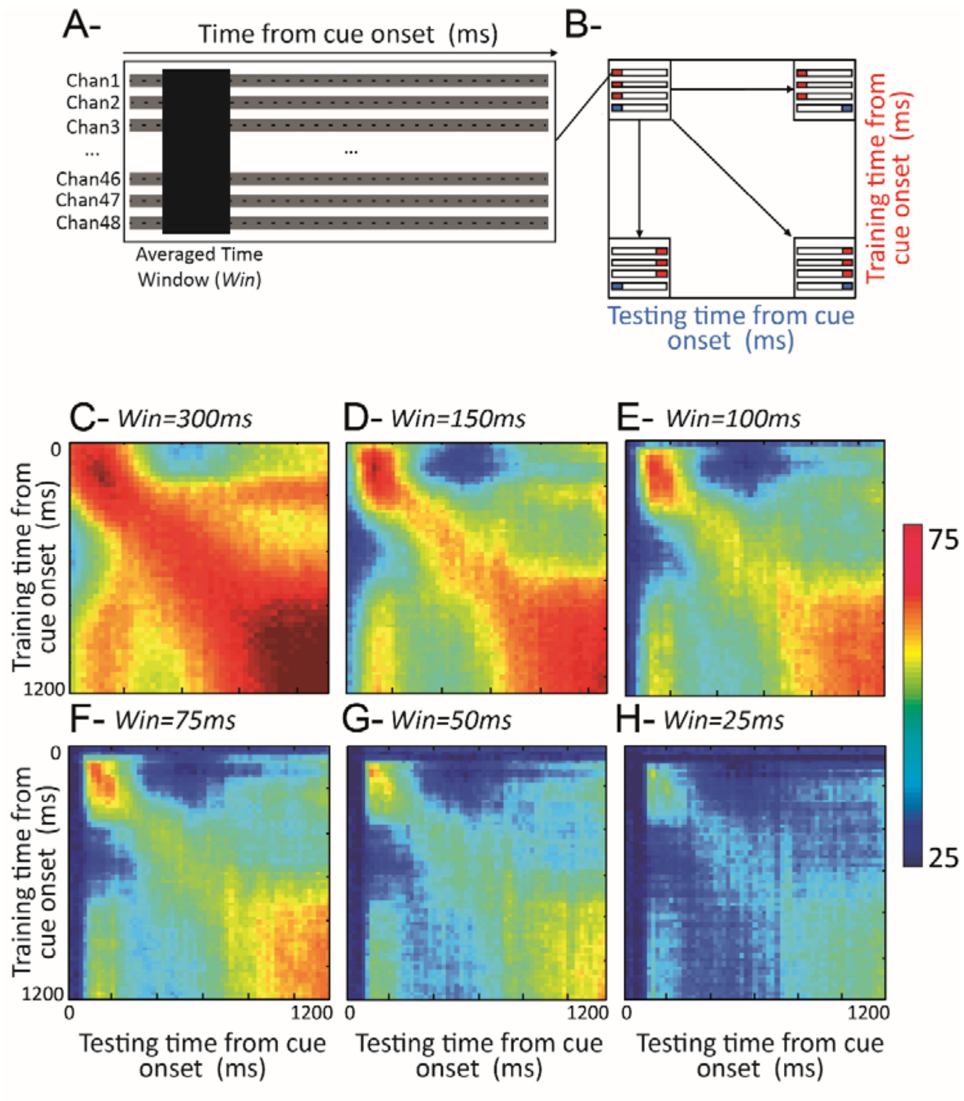


Figure S1: **Cross temporal decoding and the impact of averaging time windows.** (a) Data structure on a given trial: MUA activity is recorded onto 48 channels, in time (1 ms resolution), aligned with respect to the cue, and averaged over time windows of length *Win*. (b) Cross-temporal decoding matrices are obtained by training a decoder on activities from the 48-channels, collected at a given time *t,* averaged over ATW ms, on a subset of trials (random 70%) and testing this decoder on activities from the 48-channels, collected, averaged over ATW ms, at all possible times (resolution of 10ms), on the remaining 30% test trials. This procedure is repeated over and over by moving reference time *t* by 10 ms each time, from 0 ms to 1200 ms from cue presentation. (c-h) Cross-temporal decoding matrices with different averaging time windows from 300 ms (c), to 150 ms (d), to 100 ms (e) , to 75 ms (f) , to 50 ms (g) , to 25 ms (h) averaging window. 50 ms averaging windows reveals oscillations in the decoding performance along the testing time dimension (x-axis). These oscillations can already be seen at *Win*=75 ms
